# Exploring the Suitability of Piecewise-Linear Dynamical System Models for Cognitive Neural Dynamics

**DOI:** 10.1101/2025.01.14.633062

**Authors:** Jiemin Wu, Boateng Asamoah, Zhaodan Kong, Jochen Ditterich

## Abstract

Dynamical system models have proven useful for decoding the current brain state from neural activity. So far, neuroscience has largely relied on either linear models or nonlinear models based on artificial neural networks. Piecewise linear approximations of nonlinear dynamics have proven useful in other technical applications, providing a clear advantage over network-based models, when the dynamical system is not only supposed to be observed, but also controlled. Here we explore whether piecewise-linear dynamical system models (recurrent Switching Linear Dynamical System or rSLDS models) could be useful for modeling brain dynamics, in particular in the context of cognitive tasks. We first generate artificial neural data based on a nonlinear computational model of perceptual decision-making and demonstrate that piecewise-linear dynamics can be successfully recovered from these observations. We then demonstrate that the piecewise-linear model outperforms a linear model in terms of predicting future states of the system and associated neural activity. Finally, we apply our approach to a publicly available dataset recorded from monkeys performing perceptual decisions. Much to our surprise, the piecewise-linear model did not provide a significant advantage over a linear model for these particular data, although linear models that were estimated from different trial epochs showed qualitatively different dynamics. In summary, we present a dynamical system modeling approach that could prove useful in situations, where the brain state needs to be controlled in a closed-loop fashion, for example, in new neuromodulation applications for treating cognitive deficits. Future work will have to show under what conditions the brain dynamics are sufficiently nonlinear to warrant the use of a piecewise-linear model over a linear one.

## 1. Introduction

Applications like brain-computer/machine interfaces for controlling prostheses or responsive neuromodulation for therapeutic purposes require estimating the current brain state from neural activity [1, 2, 3, 4]. Dynamical system models improve the quality of the state estimate by reducing the problem’s dimensionality, eliminating noise, and taking advantage of knowledge about how the brain state tends to evolve over time [5, 6, 7, 8]. Dynamical system models, however, do not only allow estimating the brain state, they also support predictions of how the brain state is likely to progress in the near future, which is essential for model-based control, i.e., planning state trajectories for adjusting the brain state through stimulation [9, 10, 11, 12]. Intelligent implantable stimulators based on such a strategy could open up new avenues for the treatment of, for example, cognitive deficits resulting from neurological and psychiatric disorders [13, 14, 15, 16, 17].

Linear dynamical system models have desirable mathematical properties and seem adequate for decoding movement intentions from the motor cortex [5], but cognitive functions might be associated with more complex dynamics and require nonlinear models. Neuroscience has largely focused on artificial neural networks for learning nonlinear dynamics [18, 19, 20], but these black-box models are not ideal for model-based control. Technical applications in engineering often rely on linear approximations of nonlinear dynamics [21, 22, 23, 24, 25]. The dynamics are still considered sufficiently linear in a local neighborhood in state space, but the systems are allowed to switch/transition to a different dynamic mode based on how the state evolves. In other words, a globally nonlinear model can be sufficiently approximated by a piecewise linear model, a collection of locally linear models [26, 27]. Piecewise linear models (a particular class of hybrid systems) together with linear dynamical models are two of the main pillars of modern control [28, 29]. Here *we ask whether such piecewise linear dynamical system models could be useful for applications in neuroscience*. Dynamical system models are often estimated from continuous observations of neural activity (local field potentials or smoothed firing rates). Here *we also address the question of whether these models could be estimated from sparse firing patterns of individual neurons, as they might be observed in higher-order cortical areas related to cognitive processing*.

We approach these questions by first generating synthetic neural data based on a piecewise linear computational model of perceptual decision-making inspired by the model proposed by Wong & Wang [30]. We generate both continuous observations as well as single-neuron spike trains based on Poisson processes, whose rates are controlled by the state of the dynamical system. We demonstrate that the piecewise-linear or recurrent switching linear dynamical system (rSLDS) models [31] can be successfully recovered from the synthetic data. Furthermore, we show that the estimated rSLDS models provide a significantly better explanation for the observed data than linear dynamical system (LDS) models and, more importantly, also make significantly better predictions for future states of the system. Finally, we apply our approach to a publicly available dataset collected from nonhuman primates performing a perceptual decision task.

## 2. Method

We start by introducing the rSLDS model, explaining the theory behind it and how it is applied to neural data. We then outline the synthetic data generation process, which is essential for testing our models before applying them to actual neural data. Next, we present the evaluation metrics we use to measure the performance of the different models.

### 2.1. rSLDS model

Recurrent Switching Linear Dynamical Systems (rSLDS) [31] are an advanced class of piecewise linear dynamic models designed to decompose complex nonlinear time-series data into a series of discrete segments (modes), each governed by a simpler linear dynamical system (LDS). The rSLDS framework can be conceptualized as a composite of multiple LDS models, where transitions between these models are determined by the latent state itself. Given an rSLDS model comprising *K* distinct LDS models, or modes, and time-series data observed over *T* time steps, the model’s formulation is as follows:

For each time step *t* ∈ *T*, a *discrete* latent state *z*_*t*_ ∈ {1, 2, …, *K*} is defined to signify the LDS model currently in effect that only depends on the last state, exhibiting Markovian characteristics. This state evolves according to the logistic regression model with a weight matrix 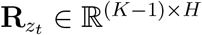 and a bias vector 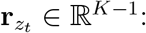

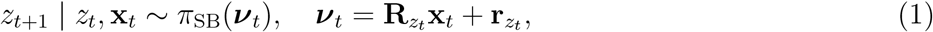

Here, **x**_*t*_ ∈ ℝ^*H*^ represents the *continuous* latent state, and *π*_SB_ : ℝ^*K−*1^ → Δ^*K−*1^ denotes a stick-breaking link function that maps the continuous state **x**_*t*_ and the discrete state *z*_*t*_ to a set of *K* normalized probabilities, governing the probabilistic transition between modes. The weight matrix 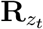 specifies the recurrent dependencies, indicating how the continuous latent state **x**_*t*_ influences the transition probabilities to the next discrete state *z*_*t*+1_. The bias vector 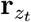 captures the Markov dependency, affecting the baseline probability of transitioning to the next discrete state *z*_*t*+1_ given the current discrete state *z*_*t*_. In this paper, we use a “recurrent only” rSLDS model [31], where all modes share the same **R** and **r**, a special case, where the next discrete state is fully determined by the current continuous state, corresponding to hard boundaries in state space between the different modes.

Upon the determination of *z*_*t*+1_, the evolution of the continuous latent state **x**_*t*+1_ is governed by:

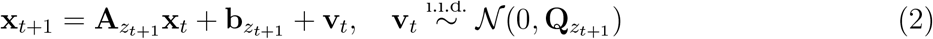

with 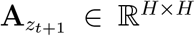 being the transition matrix, 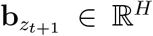 a bias vector, and **v**_*t*_ representing independent, identically distributed (i.i.d.) zero-mean Gaussian noise with covariance matrix 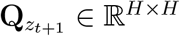 Thus, each mode is characterized by its own transition matrix **A**, bias vector **b**, and noise covariance matrix **Q**.

Subsequently, the relationship between the latent state **x**_*t*+1_ and the observed data **y**_*t*+1_ ∈ ℝ^*N*^ is established through a general linear transformation:

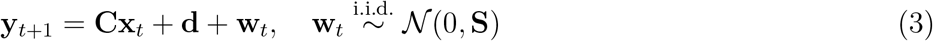

where **C** ∈ R^*N ×H*^ is the observation matrix, **d** ∈ ℝ^*N*^ is another bias vector, and **w**_*t*_ is another source of i.i.d. zero-mean Gaussian noise with covariance matrix **S** ∈ ℝ^*N ×N*^.

For modeling Poisson-distributed data **y**_*t*+1_ ∈ ℕ^*N*^, a nonlinear softplus link function is employed to ensure non-negative Poisson rates:

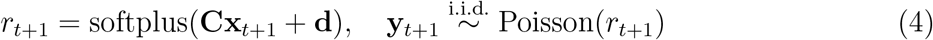

resulting in the number of spikes in each bin being drawn from a Poisson distribution with the current rate as its mean. By setting the number of modes to one (*K* = 1), the rSLDS model can be simplified to a standard LDS model. We used the Python package ssm, developed by the Linderman Lab at Stanford (https://github.com/lindermanlab/ssm) for the estimation of models, with the model fitting process relying on Variational Expectation Maximization (VEM).

When estimating hidden states using a rSLDS model, we sometimes observed discontinuities in the estimated continuous state, in particular when working with Poisson observations. Upon closer inspection, we noticed that, whenever a hidden state was to be decoded, the distribution of possible discrete states *p*(*z*_*t*_) was initialized with a uniform distribution, but had the tendency to very quickly (within just one or two iterations) settle on a particular discrete state, which constrained the continuous state to be confined to the associated part of the state space. Our solution to this issue was to modify the ssm package and to introduce a learning rate to only gradually update *p*(*z*_*t*_) and *p*(*x*_*t*_). This approach proved to be simple yet effective in mitigating the problem described above. The modified ssm code as well as additional code used for this study can be found at https://github.com/peractionlab/Decision_rSLDS.

### 2.2. Evaluation metrics

To test model quality, we used the following three metrics: (1) the evidence lower bound (ELBO), (2) the coefficient of determination (*R*^2^), and (3) the Mean Euclidean Distance (MED).

The ELBO, synonymous with the variational lower bound, serves as a metric for the log-likelihood of observational data within the framework of variational Bayesian models. It is instrumental in comparing the performance of various probabilistic models across a consistent dataset, with a larger ELBO being indicative of better model performance.

The coefficient of determination, denoted as *R*^2^, quantifies the proportion of variance in the observed data that can be explained by a model. *R*^2^ is typically used for one-dimensional, continuous data. Since we are dealing with observation vectors, we adopted the multi-dimensional *R*^2^ measure proposed by Jones & Meyer [32], which is based on the Euclidean distances between observed and predicted vectors. For multi-dimensional Poisson-distributed spike count data, we extended the deviance-based *R*^2^ measure for one-dimensional Poisson-distributed data proposed by Cameron & Windmeijer [33], assuming statistical independence between the observed spike counts for different neurons. We define the deviance term 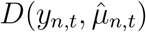 as:

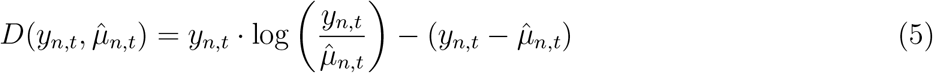

where *y*_*n,t*_ is the observed spike count for neuron *n* in a time bin at time point *t*.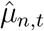 is the model’s predicted rate for the same neuron and time point. Using this deviance term, the deviance-based *R*^2^ for Poisson-distributed data is given by:

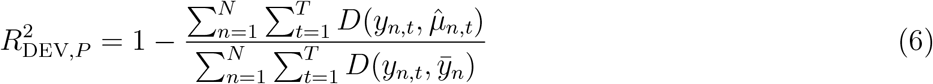

where 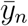 is the mean spike count for neuron *n* across all time points.

It is important to note that, while *R*^2^ can approach values close to unity in the context of assessing model quality, reflecting the extent to which measurement noise limits its value, the scenario is different for Poisson-distributed observations. Even for a model that accurately captures the underlying rates of the Poisson processes, there remains an inherent stochastic variability in the observed counts, which cannot be captured by the model and therefore imposes a ceiling on the attainable *R*^2^ values. To address this, we calculate the expected value of the numerator in the *R*^2^ computation, assuming a Poisson process with rates as determined by the model, i.e., assuming that *y*_*n,t*_ is Poisson-distributed with mean 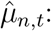

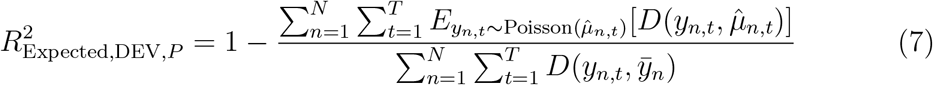

A model matching this expected *R*^2^ value can be considered an excellent model.

Finally, the mean Euclidean distance (MED) is utilized to measure the average discrepancy between the model’s predictions 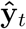 and the observed data 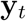, and is expressed as:

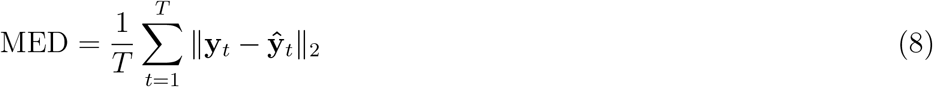

This metric directly assesses the model’s accuracy in predicting the observed phenomena.

### 2.3. Synthetic data

We first generated synthetic data on the basis of a piecewise linear adaptation of the Wong and Wang decision-making model [30], designed for decision-making between two choice options. This simulation assumed an equal provision of sensory evidence for both options on average. Since we can access the underlying dynamics, this synthetic data allowed us to test the piecewise approach.

#### Latent dynamics

The model underlying our synthetic data has a two-dimensional state space consisting of three modes (Mode 1: blue, Mode 2: red, Mode 3: yellow) as illustrated in Figure 1a. Trials initiate in Mode 1, characterized by the accumulation of noisy sensory evidence for both alternatives. This noisy evidence directs the state predominantly along the diagonal. When enough excess evidence has been accumulated in favor of one of the two choices, the system transitions into either Mode 2 or 3. These latter modes are each defined by a point attractor, which the system state converges towards. The transition from Mode 1 to the next is determined by whether the system state has crossed the boundary of the region of Mode 2 or 3. State transitions are governed by Equation 2, where *z*_*t*_ represents the active mode from the set {1, 2, 3}. Trials start at the origin (0,0) and proceed into the first quadrant, with mode switches occurring when the absolute difference between state coordinates exceeds one unit. The point attractors of Mode 2 and 3 are located at (1,6) and (6,1), respectively, and sample trajectories are depicted in Figure 1b.

**Figure 1:**
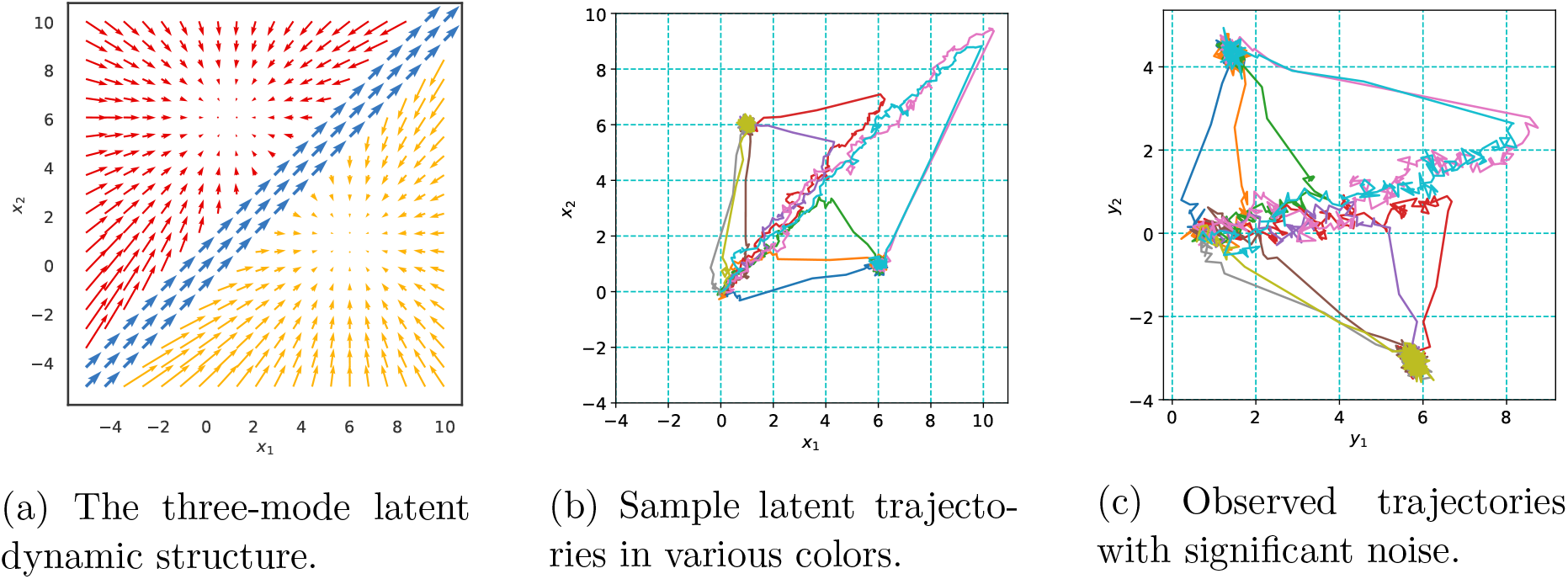
Overview of synthetic data dynamics and examples of generated trajectories.

*Observations* Employing the three-mode latent dynamic structure, we generated synthetic datasets following Gaussian and Poisson distributions, as detailed in Equations 3 and 4. Parameters *C* and *d* were adjusted to achieve a range of average firing rates. Each dataset comprised 250 trials, with 200 designated for model identification and 50 reserved for performance evaluation in a testing period. This process allowed us to ascertain the predictive capabilities of our model through a controlled assessment.

### 2.4. Visualization of higher-dimensional states

We employed Principal Component Analysis (PCA) to project these dynamics onto a 2-D embedding space for a visual representation of the inferred high-dimensional latent dynamics. PCA is a method of linear dimensionality reduction that effectively maps high-dimensional data into a lower-dimensional space, optimizing for preserving maximum data covariance and maintaining large pairwise distances.

To ensure that the choice information embedded within the inferred high-dimensional latent dynamics, denoted as *x* ∈ ℝ^*L*^, is retained as much as possible after dimension reduction via PCA, we derived the normalized difference response matrix *D* using the following computation [34]:

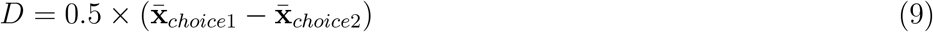

Here, 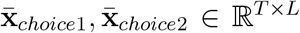 represent the trial-averaged latent dynamics. Each row within matrix *D* corresponds to the average observation difference between the two choices at a given time step. These row vectors serve as samples from which a PCA model can be fitted, yielding the final 2-D embedding of the latent dynamics, denoted as **x**_2*D*_.

### 2.5. Choice decoding

To evaluate the preservation of choice information after dimensionality reduction, we decoded the animal’s choice using a Support Vector Machine (SVM) for each time step. For each time step *t*, we combined the latent states from trajectories corresponding to choice 1 and choice 2, and then split this combined dataset into a training set (90%) and a testing set (10%). We trained an SVM classifier on the training set to classify the latent states into two classes based on their ground truth labels, performing this training for both the original 8-D latent states and the 2-D embedding latent states. After training, we applied the SVM to the testing set to predict the choice labels and computed the predictive accuracy, which is the proportion of correctly classified test samples. By comparing the predictive accuracies obtained from classifying the original 8-D latent states and the 2-D embedding, we assessed the effect of the dimensionality reduction.

### 2.6. Model comparison

To provide a clear and thorough comparison of the LDS and rSLDS models’ performance on the synthetic dataset, we designed a series of analyses focused on three critical areas: (1) estimating models, (2) making predictions, and (3) subsequent analysis of results.

#### 2.6.1. Estimating models

Prior to initiating our data analysis, we systematically partitioned the synthetic dataset into three subsets: a training set, a validation set, and a test set, adhering to a 7:1:2 ratio. Given the stochastic nature of model initialization and the utilization of the variational EM algorithm, there exists an inherent variability in the training outcomes, even when the model configurations and training data remain constant. To mitigate the risk of models converging to suboptimal solutions, particularly under conditions characterized by high noise levels and a limited dimensionality of observed variables, we implemented a strategy of training 40 instances of both the LDS and rSLDS models concurrently for each analysis setup. Subsequently, we identified the model variant that demonstrated the most robust performance, as indicated by the highest *R*^2^ value (smallest prediction error) on the validation set, for advancement to the subsequent predictive tasks and comparative result analysis.

#### 2.6.2. Making predictions

Following the training period, each model was tasked with forecasting observations (Gaussian observations or spike counts) for the next ten time points, leveraging historical data up to the current time point for every trial within the test set. These predictive outputs were then juxtaposed with their corresponding ground truth observation data and aggregated to facilitate the computation of the evaluation metrics: the coefficient of determination (*R*^2^) and Mean Euclidean Distance (MED).

#### 2.6.3. Analysis of results

The analysis of results involved a detailed examination of model performance across different hyperparameter settings, particularly focusing on the effects of varying the number of modes and latent dimensions. A Wilcoxon signed-rank test [35] was applied to the *R*^2^ and MED metrics to assess the significance of differences between the models. Additionally, we visualized the inferred latent trajectories and computed discrepancy scores relative to the ground truth model.

For the computation of discrepancy scores in relation to the ground truth model, it is essential to first map the estimated model into the same state space as the ground truth model. This alignment is necessary because, even if the estimated model converges to an optimal solution, it can still be an arbitrary linear transformation of the original model that generated the data. This arises due to the observation equation introducing a separate linear transformation between the hidden states and the observations. Therefore, before comparing the models, they must be aligned through a common linear transformation.

We denote the true model parameters as **A**_*z*_, **b**_*z*_, **C, d, R**, and **r**, and the model parameters that were estimated from the observations as **Â**_*z*_, 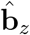, **Ĉ**, 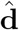, 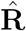, and 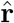. Our goal is to find the linear transformation that establishes the link between a location 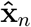 in the state space of the estimated model and the corresponding location **x**_*n*_ in the original state space:

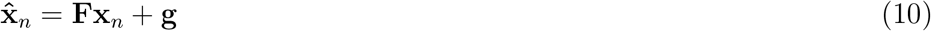

The transformation matrix **F** and the bias vector **g** can be found using the original observation equation determined by **C** and **d** and the estimated observation equation determined by **Ĉ**and 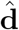, as the observations have to be the same. **F** and **g** can then be used to map the estimated model parameters back into the original state space, resulting in 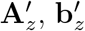, **R**^*′*^, and **r**^*′*^. **F** and **g** are determined by:

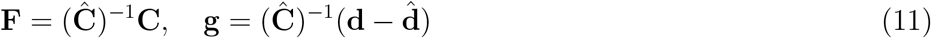

If **Ĉ**is not invertible, the Moore-Penrose pseudo-inverse can be used instead. With **F** and **g** determined, the other estimated parameters are transformed as follows:

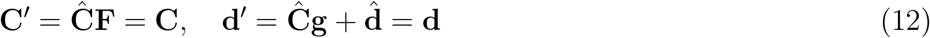

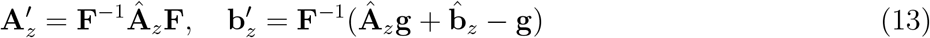

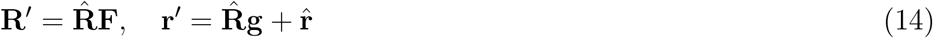

These transformations allow us to align the estimated parameters with the ground truth model for a direct comparison. Using these aligned parameters, we computed the discrepancy score, which included differences in attractor locations, switching boundaries, and eigenvalues of **A**_*z*_. Based on these scores, models were classified as “excellent” (score *<* 0.5), “good” (0.5 to 2), or “poor” (score *>* 2), providing a quantitative assessment of model fidelity to the true parameters.

The same type of transformation was also used for aligning different models in “Models for different task periods and Model initialization”.

To allow for a comparison of prediction errors across models using a Wilcoxon signed-rank test, we computed the *R*^2^ and MED distributions on a trial-by-trial basis for the test set.

## 3. Results

### 3.1. Analysis of synthetic data

#### 3.1.1. Gaussian observations

We always fitted a larger number of models of each type to the data, as variational expectation maximization does not guarantee convergence to an optimal solution. For Gaussian observations with low noise levels, we observed that a multitude of piecewise-linear model fits achieved excellent convergence, yet not all models were equally successful. The probability of identifying a superior model diminished with increasing observation noise, as depicted in Fig. 2a. Conversely, increasing the number of simultaneous observations, or the observation dimensions, significantly elevated the likelihood of discovering a model of high quality, as illustrated in Fig. 2b.

**Figure 2:**
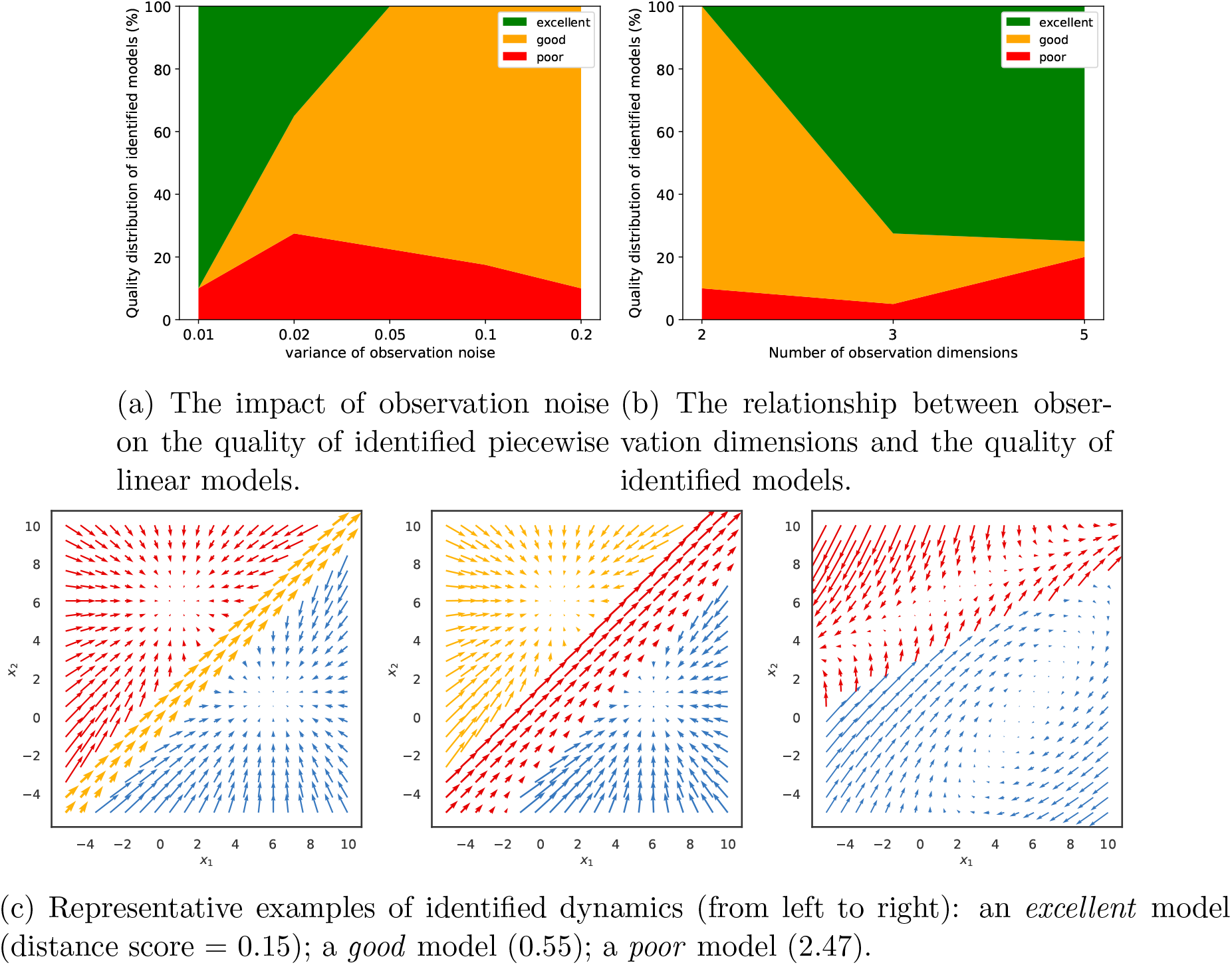
Distribution of model quality metrics across synthetic Gaussian datasets.

Building upon these findings, we categorized the trained models into three distinct tiers based on their computed distance scores relative to the ground truth dynamics: poor, good, and excellent. To provide a visual context to this classification, Fig. 2c presents the dynamic vector fields of models that correspond to each of these categories.

To test whether the rSLDS model provided a more useful description of the dynamics in terms of being able to predict how the system would evolve in the near future compared to a LDS model, we tasked both models with predicting how the (hidden) model state and the associated observations would evolve over the next 10 time steps and compared these predictions with the ground truth data. Fig. 3 elucidates the comparative performance between the LDS and rSLDS models on Gaussian data in observation space (top) and latent state space (bottom), highlighting the distinct advantages of the rSLDS model in terms of both the coefficient of determination (*R*^2^) and Mean Euclidean Distance (MED) metrics (between predicted and observed vectors). The results from a Wilcoxon signed-rank test strongly indicate a significant advantage of the rSLDS model over the LDS model (smaller prediction errors), with a p-value of less than 0.001 for all measures and all prediction horizons, underscoring the robustness of the rSLDS model in accurately capturing the dynamics of the synthetic datasets.

**Figure 3:**
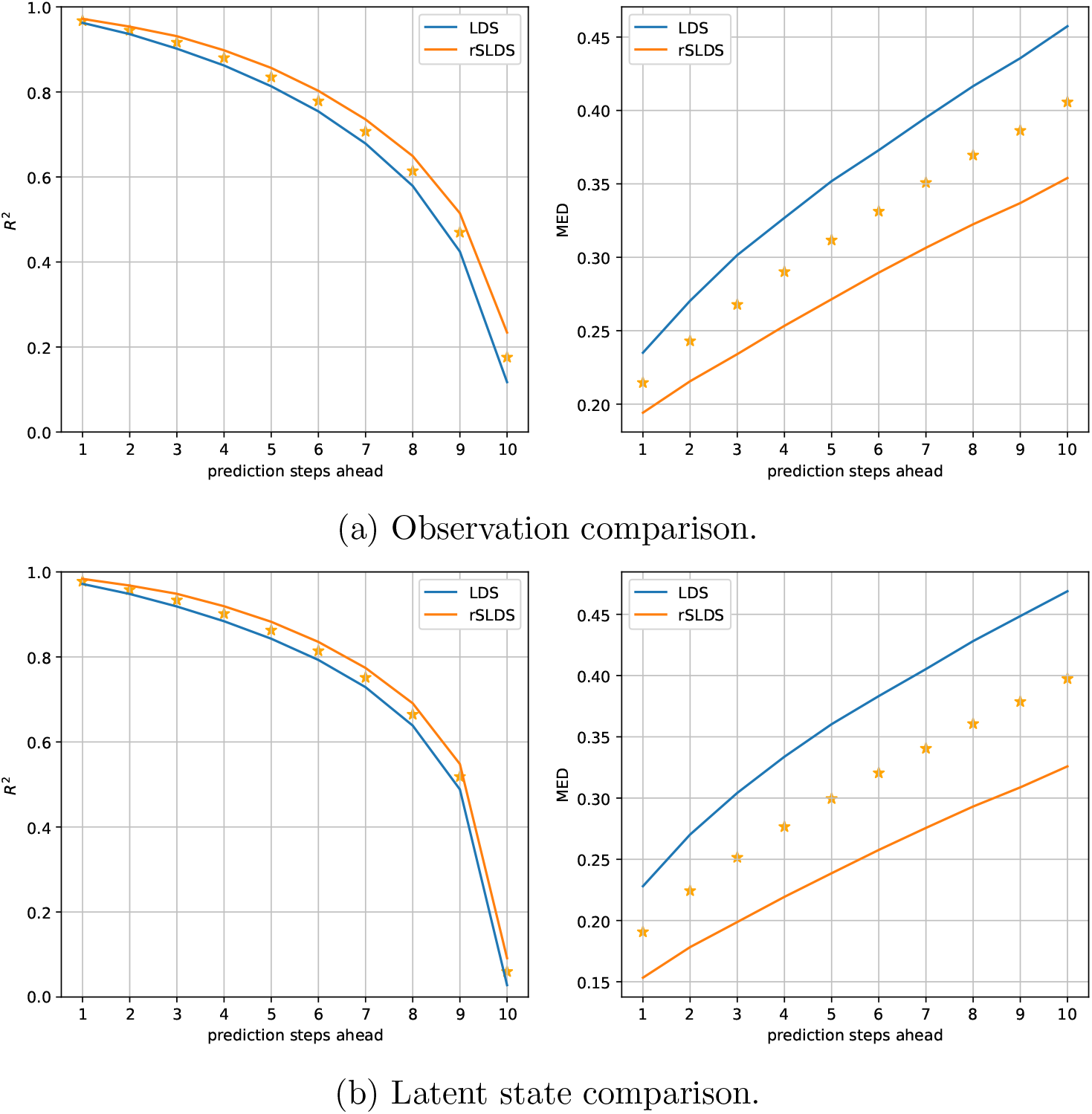
Gaussian predictive performance comparison of the LDS and rSLDS model on both the observation and latent state space, assessed using *R*^2^ and MED. The rSLDS model significantly outperformed the LDS model. The stars indicate a significant difference between both models at p*<*0.001.

#### 3.1.2. Poisson observations

When dynamical system models have to be estimated from single-unit spiking data, the typical approach is to count spikes in bins of some chosen width, apply some smoothing, and treat the result as continuous observations. However, when dealing with neurons with low firing rates, like in the case of the prefrontal dataset we will be looking at later, with average firing rates of about 10 spikes/s, the spike count data can become pretty sparse (with a substantial number of zero counts), and this approach doesn’t work anymore. We therefore wanted to test whether it is still possible to estimate rSLDS models from these data, treating them as Poisson rather than continuous observations. Since the variance of a Poisson distribution is identical to its mean, the inherent observation noise will typically be larger than what we have been considering in the case of Gaussian observations.

As a consequence, as can be seen in Fig. 4a, the number of excellent fitted models is much smaller. The quality of the fits depends on the expected number of spikes per bin and on the number of observed neurons. The higher the Poisson rate, the more likely it is to obtain a good or perhaps even excellent model. About 20% of the fitted models were still good for an expected spike count of 0.5 spikes per bin. We would not recommend applying the technique to much lower rates. For an expected spike count of 2 spikes per bin, a few excellent models were only observed for a large number of observations (see Fig. 4b). At the same time, the number of poor models also increased with the number of observations (neurons), likely due to the increasing number of model parameters, which increases the risk of getting stuck in suboptimal solutions. As mentioned before, it is therefore important to always fit a larger number of models and to select the best one.

**Figure 4:**
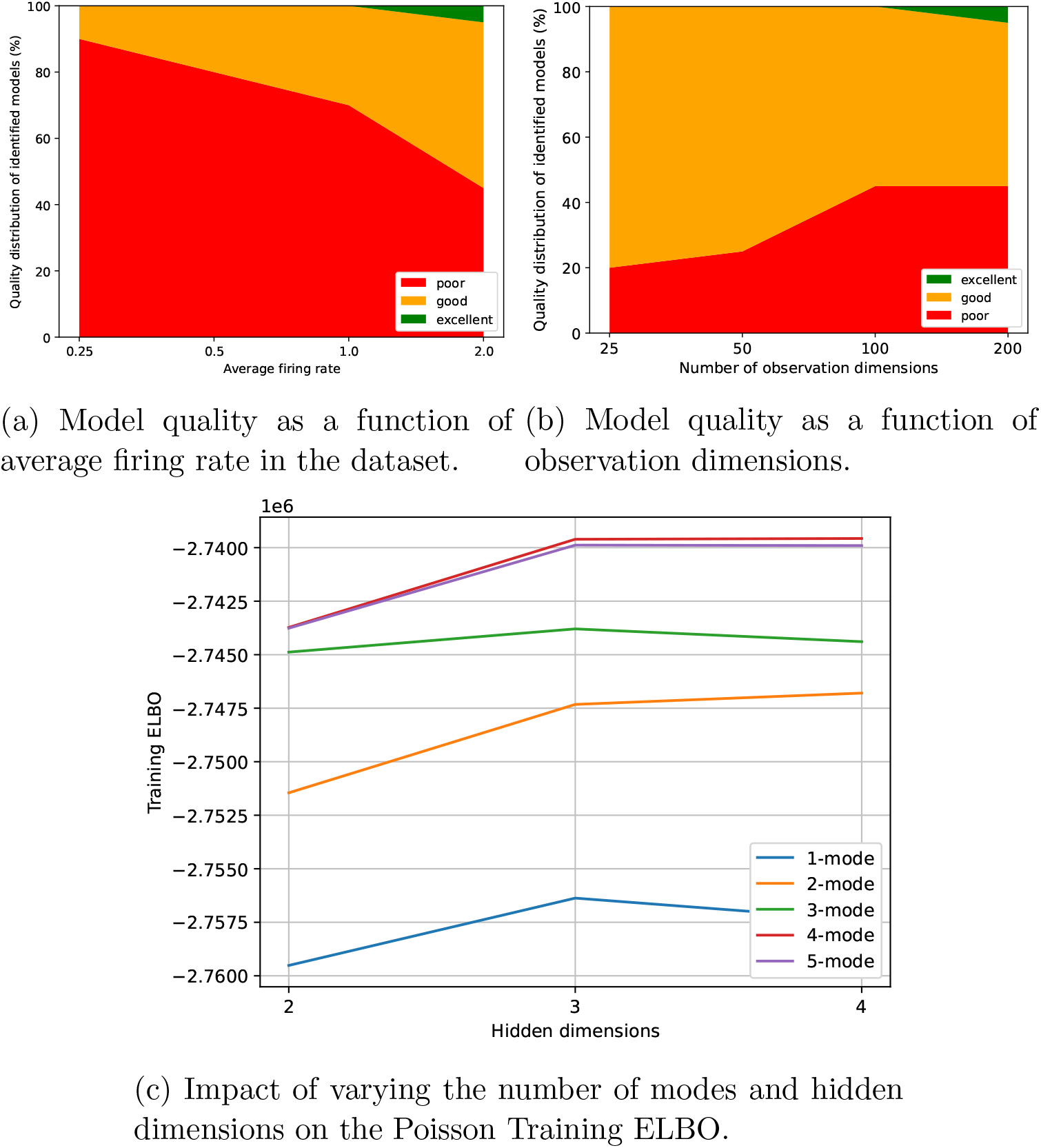
Distribution of model quality metrics across synthetic Poisson datasets.

When creating our synthetic dataset, we used a model with two hidden dimensions and three modes. When fitting models with various numbers of hidden dimensions and modes to the data, the Evidence Lower Bound (ELBO) indicated that, when starting with a 2-dimensional model with one mode, adding a 2nd and a 3rd mode to the model improved the quality of the fit much more than adding more hidden dimensions (see Fig. 4c).

Despite the larger challenge of estimating a good rSLDS model from the Poisson data compared to the Gaussian data earlier, its predictive accuracy still surpasses that of an LDS model, as illustrated in Fig. 5.

**Figure 5:**
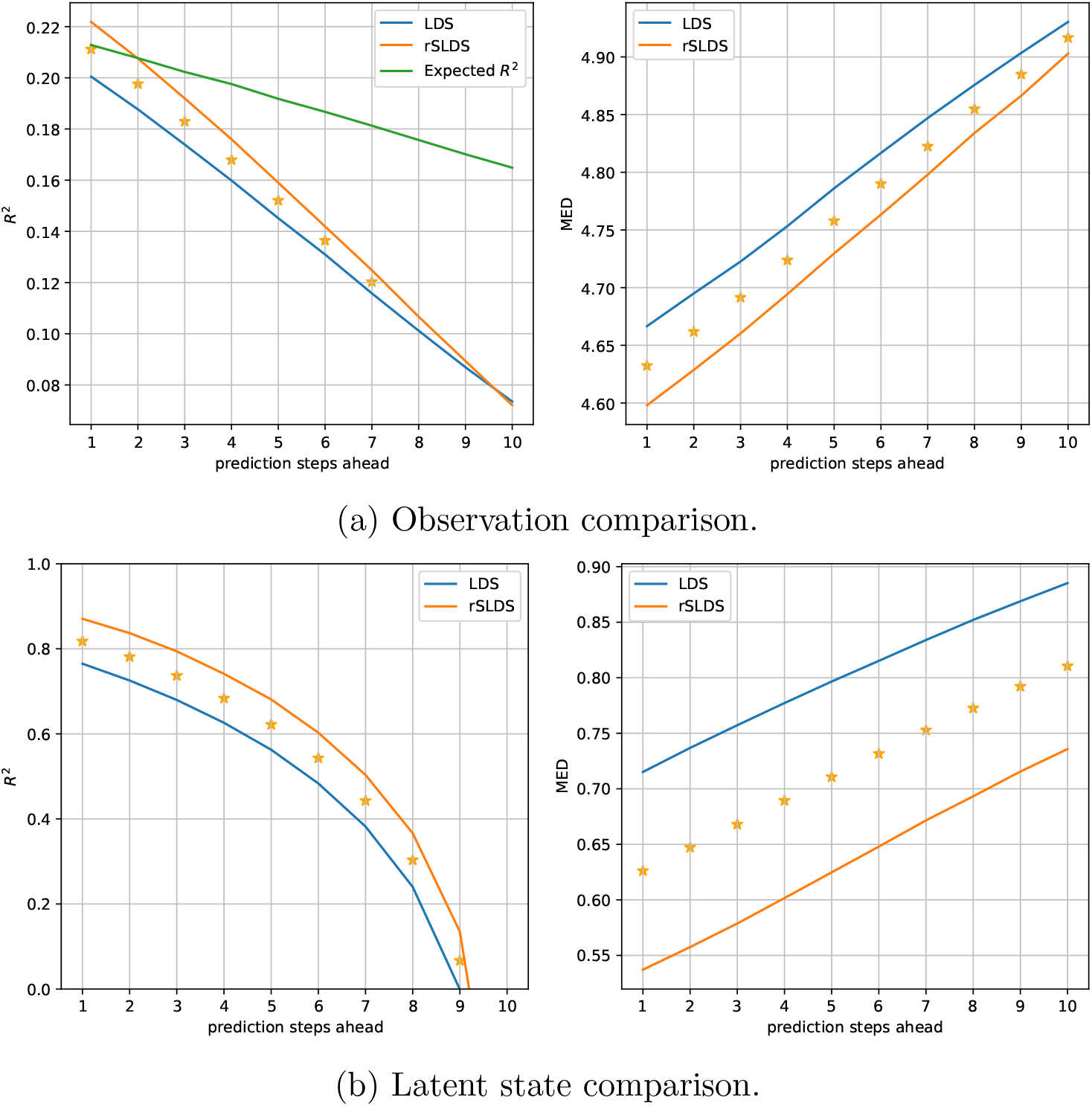
Poisson predictive performance comparison of the LDS and rSLDS model on both the observation and latent state space, assessed using *R*^2^ and MED.

An analysis leveraging the coefficient of determination (*R*^2^) and Mean Euclidean Distance (MED) metrics demonstrates that the rSLDS model significantly outperforms the LDS model in forecasting up to at least 7 time steps ahead. Due to the inherent variability of the Poisson observations, the observed *R*^2^ values are much lower than in the earlier case of Gaussian observations. To still get an idea of the quality of the estimated models, we calculated what the *R*^2^ value was expected to be, if the model had perfectly captured the Poisson rates, which we refer to as the “Expected *R*^2^” (see Section 2), plotted in green in Fig. 5a. The results suggest that our rSLDS model is close to optimal for predictions one and two time steps ahead.

Since we know the ground-truth hidden states underlying our synthetic data, we can also calculate the predicition error in the latent state space, which is shown in Fig. 5b. Since this eliminates the inherent variability of the Poisson observations, the observed *R*^2^ values are much larger, and the significant advantage of the rSLDS model compared to the LDS model is more obvious and statistically significant for all analyzed prediction horizons. This indicates that the rSLDS model outperforms the LDS model in terms of decoding and predicting system states from Poisson observations. We will not be able to perform this latter analysis for the real neural data, as the ground-truth hidden states are unknown.

### 3.2. Analysis of real neural data from perceptual decision-making

To test our approach with real neural data, we took advantage of a publicly available dataset of approx. 200 simultaneously recorded neurons in prefrontal cortex of monkeys performing a perceptual decision task. The “Kiani dataset” was derived from a direction discrimination task involving macaque monkeys trained to perform a task, in which they viewed a patch of randomly moving dots for 800 ms, followed by a variable-length delay period [36]. At the end of the delay, a “Go” cue prompted the monkeys to report their perceived motion direction by making a saccadic eye movement to one of two targets (T1 or T2). We analyzed either data from the whole trial or from two particular time periods: the “Decision Period”, which extends from the onset of the dots to the “Go” cue, representing the time during which the monkey makes its decision, and the “Motor Period”, which was defined as starting 100 ms before the “Go” cue and ending 150 ms after the saccade onset, indicating the window in which the monkey prepares and executes its eye movement response. Neural activity was simultaneously recorded from hundreds of single- and multi-neuron units in the prearcuate gyrus (area 8Ar) using a 96-channel multi-electrode array, which covered a 4 mm × 4 mm area of the cortical surface. This setup allowed for the dynamic decoding of the decision variable and the prediction of the monkeys’ choices based on neural population responses.

In this section, we delve into the analysis of the Kiani dataset, aiming to uncover insights into the neural dynamics underlying the Decision and Motor Periods. We begin by comparing the predictive performance of two models, the linear dynamical system (LDS) and the recurrent switching linear dynamical system (rSLDS), across varying levels of model complexity, using data from the full trials. We then explore the Decision and Motor Periods separately to address whether the underlying dynamics are different or the same. Subsequently, we examine how well the animal’s choice can be decoded from estimated hidden states.

#### 3.2.1. LDS versus rSLDS models

The number of hidden dimensions is a pivotal hyperparameter for both linear and piecewise linear models in modeling neural data. To evaluate the efficacy of these models in capturing the latent dynamics encoded in the neural data, we conducted a comparative analysis using the T33 session from the Kiani dataset, the session with the best behavioral performance. The dataset was segmented into 80 ms-long time bins, a duration chosen based on the average firing rate of the recorded neurons, which is approximately 10 spikes per second. This results in an expected spike count per bin of about 0.8, a value for which we anticipate being able to estimate good models, based on our insights from the synthetic data.

We estimated models with hidden dimensions varying from 2 to 24, aiming to predict the spike count of the next time bin based on the current one. The predictive accuracy of these models was again measured using the coefficient of determination (*R*^2^) and Mean Euclidean Distance (MED) metrics. As illustrated in Fig. 6, the comparative results show that the performance metrics for both LDS and rSLDS models improve with an increasing number of hidden dimensions. Notably, as the number of hidden dimensions increases, the *R*^2^ values approach the expected value for Poisson-distributed observations with the rates predicted by the model, indicating that our largest models (24 hidden dimensions) capture almost as much variability as they possibly can for Poisson observations.

**Figure 6:**
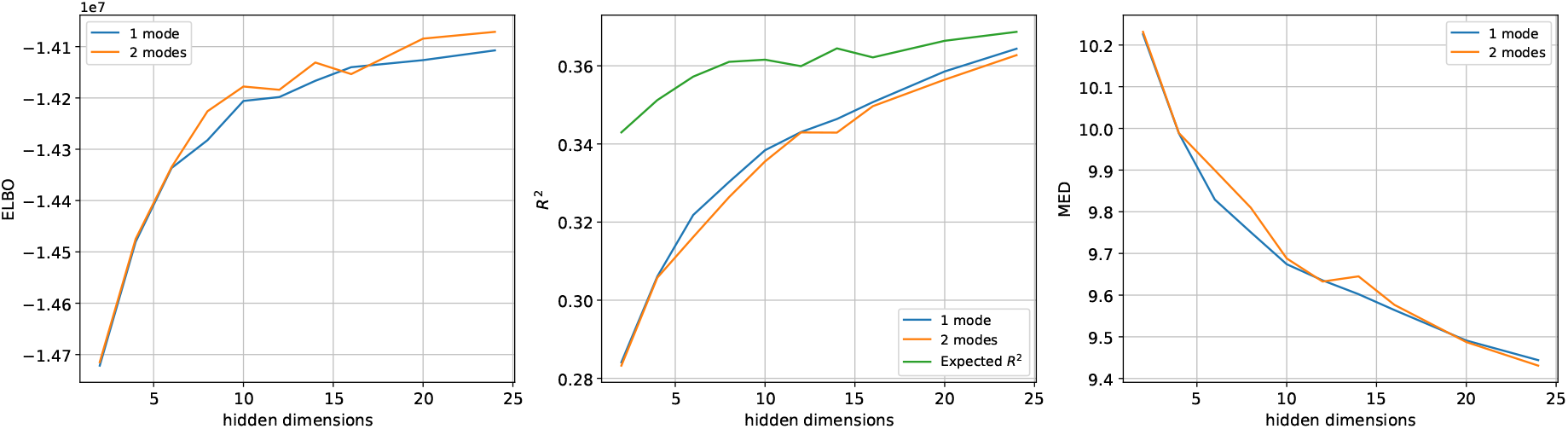
Performance comparison of 1-mode LDS and 2-mode rSLDS models across a range of hidden dimensions on the T33 dataset. The performance of both models escalates with additional hidden dimensions and eventually tends to plateau. No distinct superiority of the rSLDS model is observed across all three evaluation metrics when juxtaposed with the LDS model.

Our results on synthetic data, which were inspired by a popular nonlinear computational model of perceptual decision-making, have shown that the rSLDS model was able to capture these nonlinearities and provide a better explanation for the data than a linear model. This led us to expect that the rSLDS model should also achieve superior predictive performance over the LDS model for the Kiani dataset. However, contrary to these expectations, rSLDS models with two modes did not significantly exceed the LDS models in performance (see Fig. 6). The same qualitative observation was made when using data from the C42 session and for 40 ms-long time bins (data not shown). To rule out that the missing benefit of additional modes was not just due to the rSLDS models not having converged to an optimal solution, we conducted further analyses.

#### 3.2.2. Models for different task periods and Model initialization

First, we fitted separate linear models based on data from the Decision Period and the Motor Period. To be able to visualize the results, we use models with two hidden dimensions. The resulting vector fields, plotted in a common state space (see Section 2) are shown on the left side of Fig. 7. The dynamics during the decision period were found to be stable, while those in the motor period were characterized by instability. To maximize the chances of a 2-mode rSLDS model being able to capture this difference in dynamics for different task periods, we initialized the two modes with these two linear models. The boundary between the two modes was determined as follows: we calculated the latent trajectories for all trials during different task periods and identified the midpoints of these trajectories for each phase. The perpendicular bisector of the line segment connecting these two midpoints was then used as the boundary between the two modes (see middle panel of Fig. 7).

**Figure 7:**
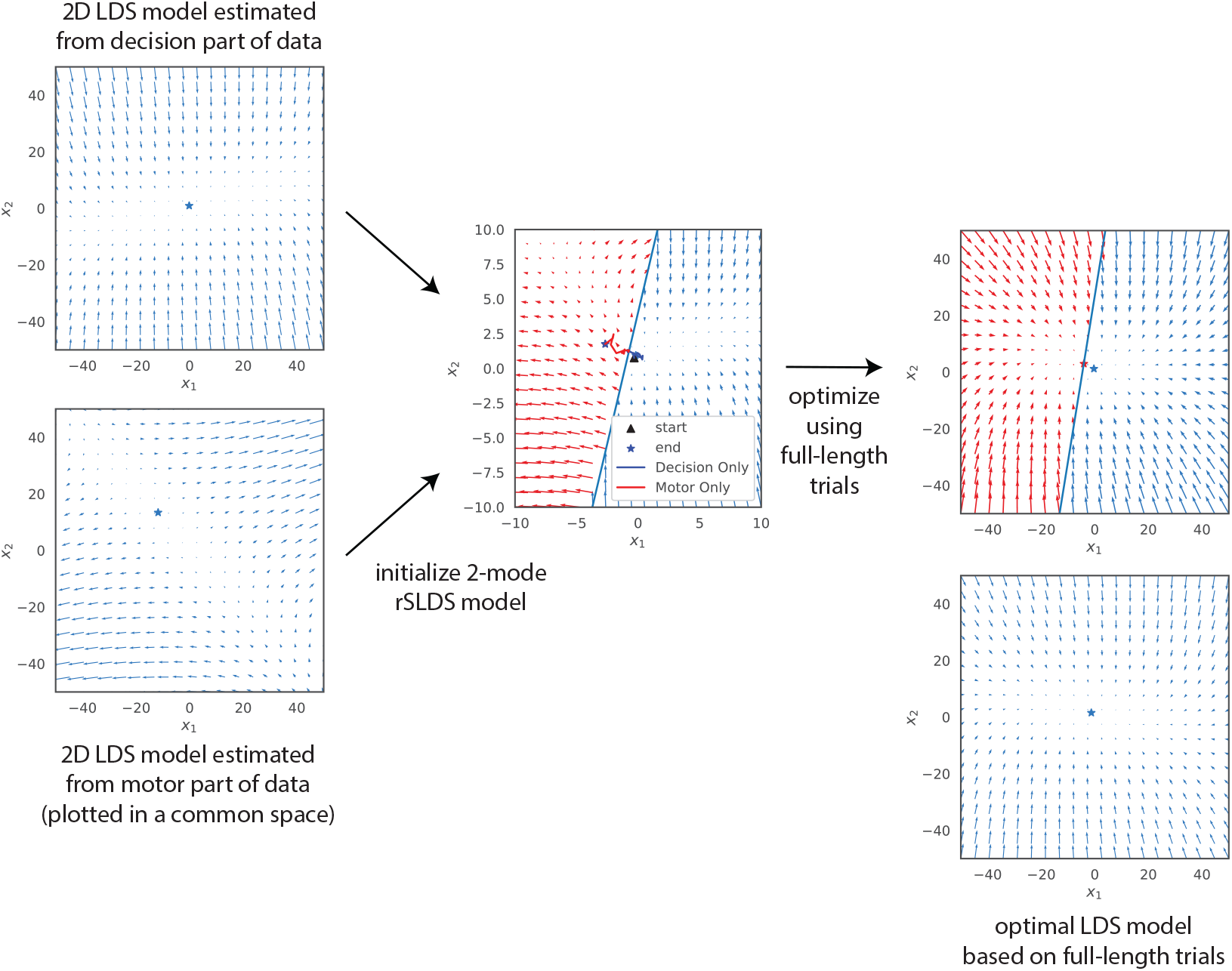
Inferred latent dynamics from an LDS model and a 2-mode rSLDS model, initialized by integrating dynamics from separate LDS models optimized on the decision and motor period datasets. Despite this initialization that should maximize the chance of finding a piecewise model that combines the decision and motor dynamics, both models converge to a similar latent dynamic following training.

The result after optimizing the model using the full trial data is shown in the top-right panel of Fig. 7. The unstable dynamics have disappeared, and the result is virtually identical to a linear model fitted to the same data (bottom-right panel), again plotted in a common state space.

Considering that the Decision Period is substantially longer than the Motor Period, we questioned whether the Variational EM algorithm might favor a model that more accurately represents the decision period, potentially neglecting the motor period due to its relatively minor contribution in terms of data points. To address this, we constructed a dataset containing only the final part of the Decision Period and the entire Motor Period, equalizing their durations. Despite these modifications, our findings mirrored those presented in Fig. 7. Consequently, we conclude that a piecewise linear model with a common observation equation cannot capture the observed change in dynamics from the decision period to the motor period. We will revisit this observation later in Discussion.

#### 3.2.3. Decoding accuracy

Since the neural activity was recorded while monkeys were making perceptual decisions, we also wanted to assess how well an upcoming choice could be decoded from estimated hidden states, depending on which dynamical system model was used. Fig. 6 had shown that adding hidden dimensions (within the studied range of up to 24 dimensions) continually increased the model’s ability to capture and predict the neural activity. This prompted the question how the ability to decode the choice would depend on the dimensionality of the hidden state space. For each of the models, we first decoded the latent trajectories associated with each experimental trial. Subsequently, for each time step, we trained a Support Vector Machine (SVM) algorithm [37] on the training dataset. After establishing the SVM model, we evaluated its performance on a separate test set by measuring the accuracy of choice classification as a function of time.

Fig. 8 illustrates that the rSLDS model’s choice-decoding accuracy progressively increases across a broad range of hidden dimensions, highlighting the dependency of choice information preservation on the model’s dimensionality. With a limited number of hidden dimensions (*H*), such as 2 to 4, the decoding accuracy remains limited and only deviates from chance towards the end of the decision trial, when the animal reports its choice, long after the motion viewing period. For 6 or 8 dimensions, the ability to decode choice starts earlier in the trial. Finally, for 10 dimensions or more an accuracy of about 80% is reached towards the end of the motion viewing period, which then further increases and approaches 100% towards the end of the trial.

**Figure 8:**
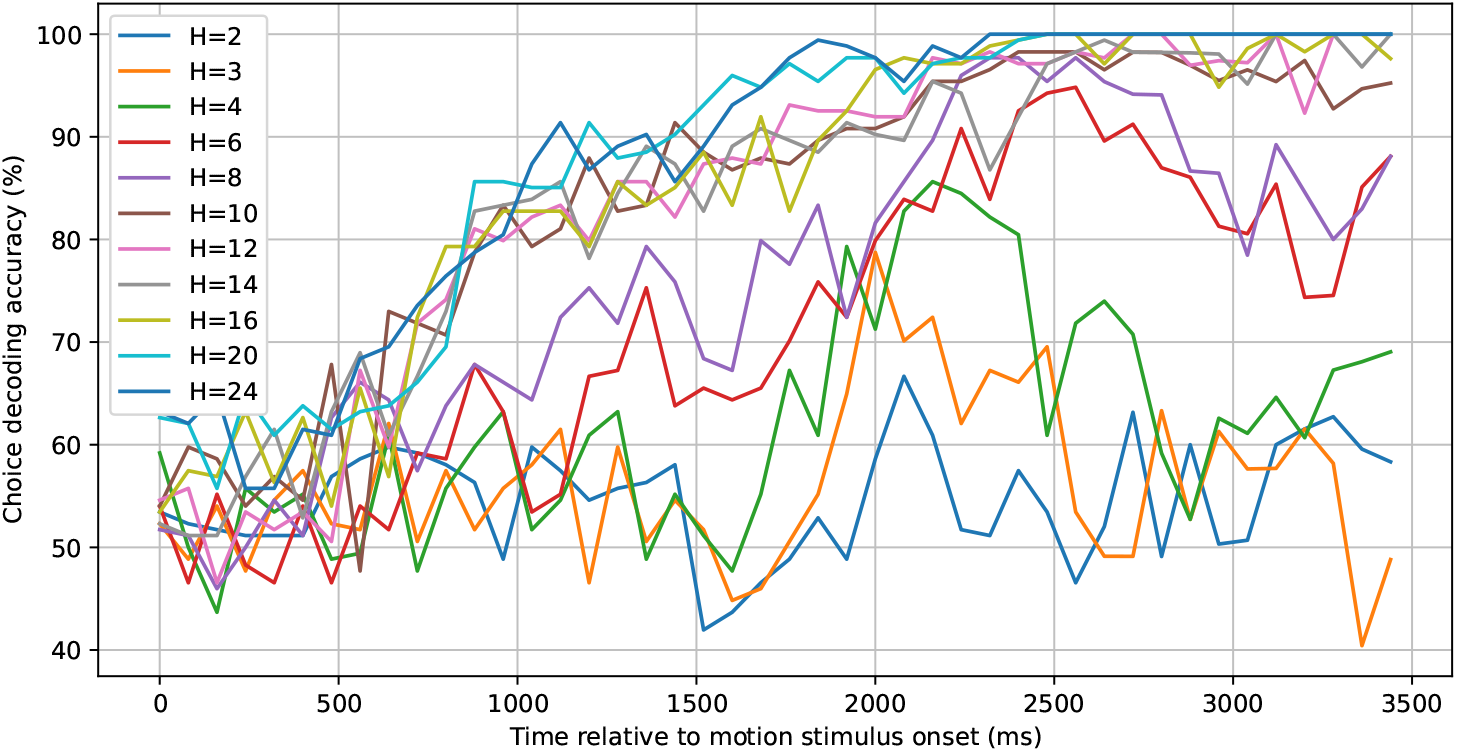
Choice decoding accuracy based on inferred latent states for different numbers of hidden dimensions. A linear SVM decoder is applied to the latent states from various trials to ascertain the corresponding choice label at each time step. The decoding accuracy surpasses 90% approx. 2000 ms after the onset of the random dot motion stimulus for models with at least 10 hidden dimensions.

#### 3.2.4. High-dimensional dynamics embedding

While an expanded number of hidden dimensions enhances choice decoding accuracy and augments the model’s predictive capabilities, it introduces challenges in visualizing the dynamics. To address this issue, we started with an 8-dimensional model and determined a two-dimensional projection of the hidden states that would maximize the separation between the states associated with both possible choices (see Section 2 for details). Fig. 9a presents the average latent trajectories in this two-dimensional space for both possible choices. The trajectories start at the same location, but diverge as time progresses, and end at very different locations. To assess the efficacy of this 2D embedding approach, we recomputed the choice decoding accuracy on the embedded trajectories. The results, depicted in Fig. 9b, closely resemble those derived from the original 8-dimensional trajectories, indicating that almost all choice information is contained in a particular two-dimensional subspace.

**Figure 9:**
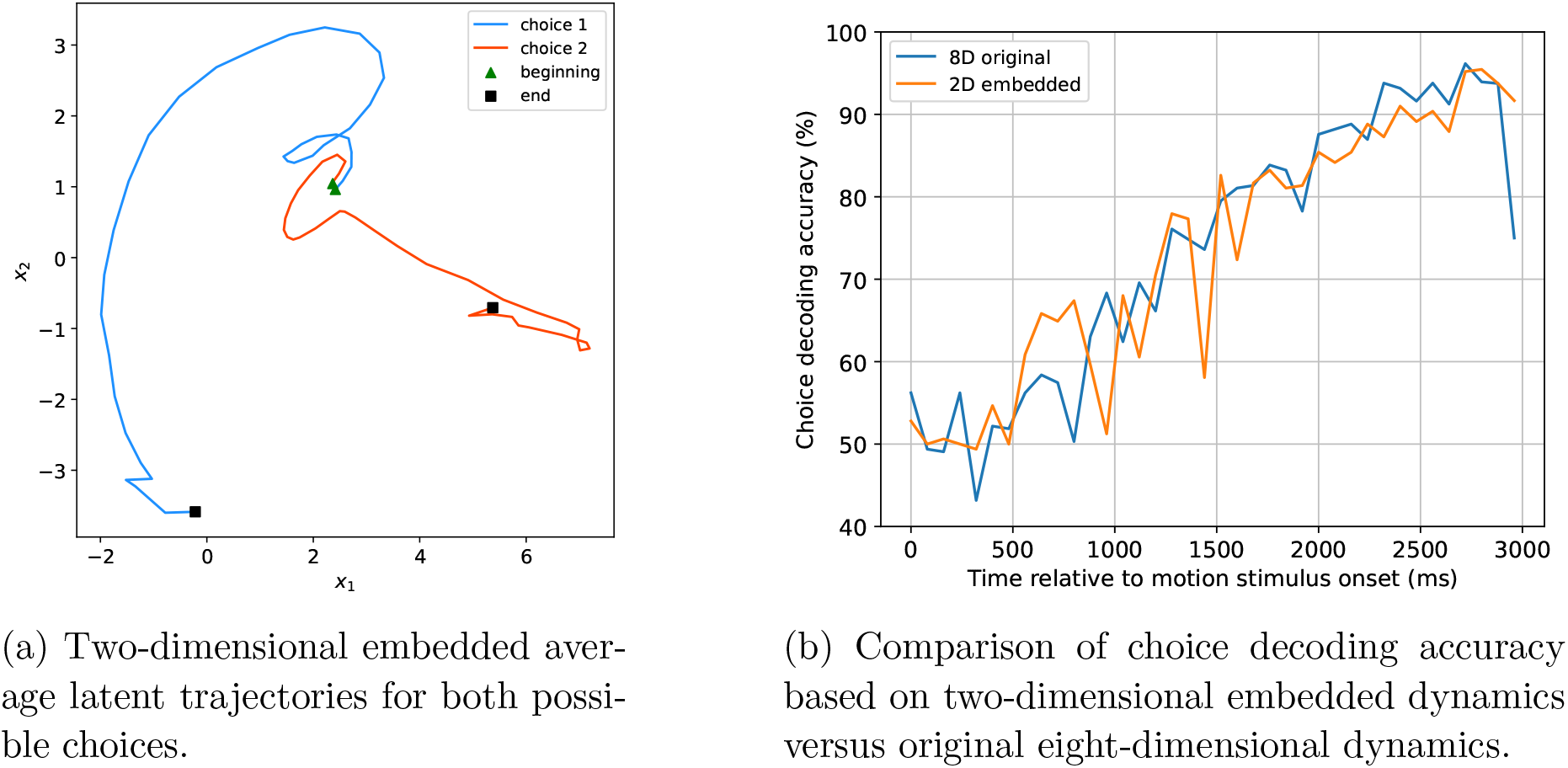
2D projection approach maintaining choice decodability

## 4. Discussion

### 4.1. Successful estimation of piecewise-linear models from synthetic neural data and superior prediction performance

Using synthetic neural data generated by a piecewise-linear adaptation of the computational model proposed by Wong & Wang [30] for perceptual decision-making, we were able to demonstrate that the nonlinear structure of the dynamical system could be successfully estimated from the data. Not surprisingly, the quality of the estimated models depended on how many signals were observed and the amount of noise on these observations. For continuous observations, the estimated models tended to be of good quality, as long as the number of observed signals matched at least the dimensionality of the hidden state space and the observation noise was not too large. Estimating the model structure from Poisson-distributed single-unit spiking data was more challenging, but still possible. These observations are inherently more noisy, due to the stochastic nature of the Poisson process, and potentially quite sparse, if the firing rates of the observed neurons are rather low. Good models could still be obtained as long as the number of observed neurons was at least an order of magnitude larger than the dimensionality of the hidden state space and the average expected spike count per time bin wasn’t much lower than 0.5. Since the model estimation based on Variational Expectation Maximization can get stuck in suboptimal solutions, we recommend fitting a larger number of models and selecting the best one.

Perhaps more importantly, the estimated piecewise-linear dynamical models were able to make significantly more accurate predictions for how the state of the system would evolve in the near future than just a linear dynamical system model. This could give these piecewise-linear models a significant advantage when considering applications like closed-loop neural modulation, where state predictions can be used for model-based control.

### 4.2. Piecewise-linear models did not outperform their linear counterparts when applying the approach to a publicly available dataset recorded from prefrontal cortex of monkeys performing perceptual decisions

To our surprise, we did not see the same benefit when applying our approach to a publicly available dataset of about 200 simultaneously recorded neurons in prefrontal cortex of monkeys performing perceptual decisions. Standard linear dynamical system models and piecewise-linear models provided virtually identical ability to explain the neural data and prediction performance. This does not imply that the same would have to be true for other datasets obtained from other brain areas or when performing different tasks, but it certainly warrants some discussion.

One possibility would be that, for some reason, the structure of the dynamical system, although compatible with our piecewise-linear model, could not be estimated from the available data. Given that the number of available neurons was at least an order of magnitude larger than the dimensionality of the hidden state space and given that we chose a time bin width of 80 ms, which resulted in an average expected spike count per bin of 0.8, a value that allowed robust model estimation when using our synthetic data, this seems unlikely.

Another possibility would be that the dynamical system, although nonlinear, is incompatible with certain assumptions made by our model. We will return to this topic in our discussion further below.

Finally, it could be that the neural dynamics are not sufficiently nonlinear to warrant fitting a nonlinear model. Some recent observations in the literature provide some support for this idea. For example, Nozari et al. [6] analyzed resting-state neural activity (both fMRI and intracranial EEG data) and fitted a wide range of both linear and nonlinear dynamical models. The model best capturing the data turned out to be a linear autoregressive model, suggesting that the dynamics were not sufficiently nonlinear to give a nonlinear model an advantage. Galgali et al. [34] analyzed the same dataset by Kiani et al. that was also used for our analysis here and made a very similar observation: The neural dynamics during a particular task period, like the decision formation period and the saccadic eye movement period, were well-described by a linear dynamical model.

This raises the final question why the different linear dynamics during the different task periods could not be captured by a piecewise linear model with multiple modes. The rSLDS model we have been using is designed to approximate one larger nonlinear flow field through breaking it down into multiple locally linear modes or regions in state space. There is still one common observation equation that applies to all modes/regions. The model switches to a different mode due to the location in state space changing, and the observations (firing pattern of the neurons) have to change accordingly. In contrast, the change in linear dynamics when transitioning between different task periods might happen without a major change in the firing pattern/location in state space and might therefore be better described by a time-variant linear system rather than a nonlinear system. Future research will have to show whether time-variant models can provide a better description of the neural dynamics.

## 5. Conclusion

We have set out to evaluate the suitability of piecewise-linear dynamical system models for capturing cognitive neural dynamics. We were able to demonstrate that piecewise-linear dynamical system (rSLDS) models could be successfully estimated from synthetic neural data generated by a nonlinear computational model of perceptual decision-making. Importantly, the rSLDS model significantly outperformed a standard LDS model in terms of predicting the state of the dynamical system in the near future. Given the straightforward mathematical structure of a piecewise-linear dynamical system model, it would be predestined to be used for model-based control applications, e.g., closed-loop neuromodulation for the treatment of cognitive deficits resulting from neurological or psychiatric disorders, when nonlinearities of the dynamics need to be captured.

Much to our surprise, when applying our modeling approach to a publicly available dataset of approximately 200 simultaneously recorded neurons in prefrontal cortex of monkeys performing perceptual decisions, we did not find a significant advantage of rSLDS models over standard LDS models, suggesting that the neural dynamics were not sufficiently nonlinear at any given time to warrant the use of a nonlinear model. Future research will have to show whether this finding applies to a wide range of cognitive neural datasets, including different types of neural activity (single-unit spiking, multi-unit activity, LFPs, iEEG) and recordings from different brain areas and during different tasks, or whether it was due to specific features of the particular dataset that was used for our analysis, and other datasets will show a benefit of using piecewise-linear dynamic models, similar to our synthetic data.

## Acknowledgments

This work was partially supported by the National Science Foundation (grant number 2024526).

## Notes

### Competing Interest Statement

The authors have declared no competing interest.

